# SEG: Segmentation Evaluation in absence of Ground truth labels

**DOI:** 10.1101/2023.02.23.529809

**Authors:** Zachary Sims, Luke Strgar, Dharani Thirumalaisamy, Robert Heussner, Guillaume Thibault, Young Hwan Chang

## Abstract

Identifying individual cells or nuclei is often the first step in the analysis of multiplex tissue imaging (MTI) data. Recent efforts to produce plug-and-play, end-to-end MTI analysis tools such as MCMICRO^1^– though groundbreaking in their usability and extensibility – are often unable to provide users guidance regarding the most appropriate models for their segmentation task among an endless proliferation of novel segmentation methods. Unfortunately, evaluating segmentation results on a user’s dataset without ground truth labels is either purely subjective or eventually amounts to the task of performing the original, time-intensive annotation. As a consequence, researchers rely on models pre-trained on other large datasets for their unique tasks. Here, we propose a methodological approach for evaluating MTI nuclei segmentation methods in absence of ground truth labels by scoring relatively to a larger ensemble of segmentations. To avoid potential sensitivity to collective bias from the ensemble approach, we refine the ensemble via weighted average across segmentation methods, which we derive from a systematic model ablation study. First, we demonstrate a proof-of-concept and the feasibility of the proposed approach to evaluate segmentation performance in a small dataset with ground truth annotation. To validate the ensemble and demonstrate the importance of our method-specific weighting, we compare the ensemble’s detection and pixel-level predictions – derived without supervision - with the data’s ground truth labels. Second, we apply the methodology to an unlabeled larger tissue microarray (TMA) dataset, which includes a diverse set of breast cancer phenotypes, and provides decision guidelines for the general user to more easily choose the most suitable segmentation methods for their own dataset by systematically evaluating the performance of individual segmentation approaches in the entire dataset.

## INTRODUCTION

Highly multiplexed tissue imaging (MTI) techniques allow visualization and quantification of the spatially resolved expression of various protein markers at single-cell resolution in tissues^2–4^. MTI downstream analyses including individual cell phenotyping, cell population, and spatial analyses rely on cell and nuclei segmentation^5^. As segmentation is the downstream analyses cornerstone, it is necessary to evaluate and choose the most appropriate segmentation methods. Deep learning-based methods specifically pre-trained for biomedical image segmentation are considered state-of-the-art and can provide good out-of-the-box results^6–12^. However, the published performance of the pre-trained models may be untrustworthy on the user’s data or new types of data, especially when test images are very different from the training images because of the variability of biological samples.

As shown in Figure 1A for the analysis of human tumor atlas network (HTAN) TNP-TMA core G5 (see details Dataset in METHODS section), we often observe that segmentation results from different methods show inconsistencies and discrepancies in segmentation tasks and thus cause feature level differences; for example, 1) the number of segmented nuclei ranges widely, from the lowest number of 6,662 using Stardist^6^ to the highest number of 23,600 using UnMICST^12^, and 2) further, the Cellpose^3^- and UnMICST-based mean intensity histograms in core F4 show three distinct peaks for the pan-cytokeratin marker while only two are observed for other methods. As inaccurate segmentation introduces systematic error or bias in downstream analysis, it is important to evaluate segmentation methods and choose the most suitable one for a specific application.

**Figure 1.**
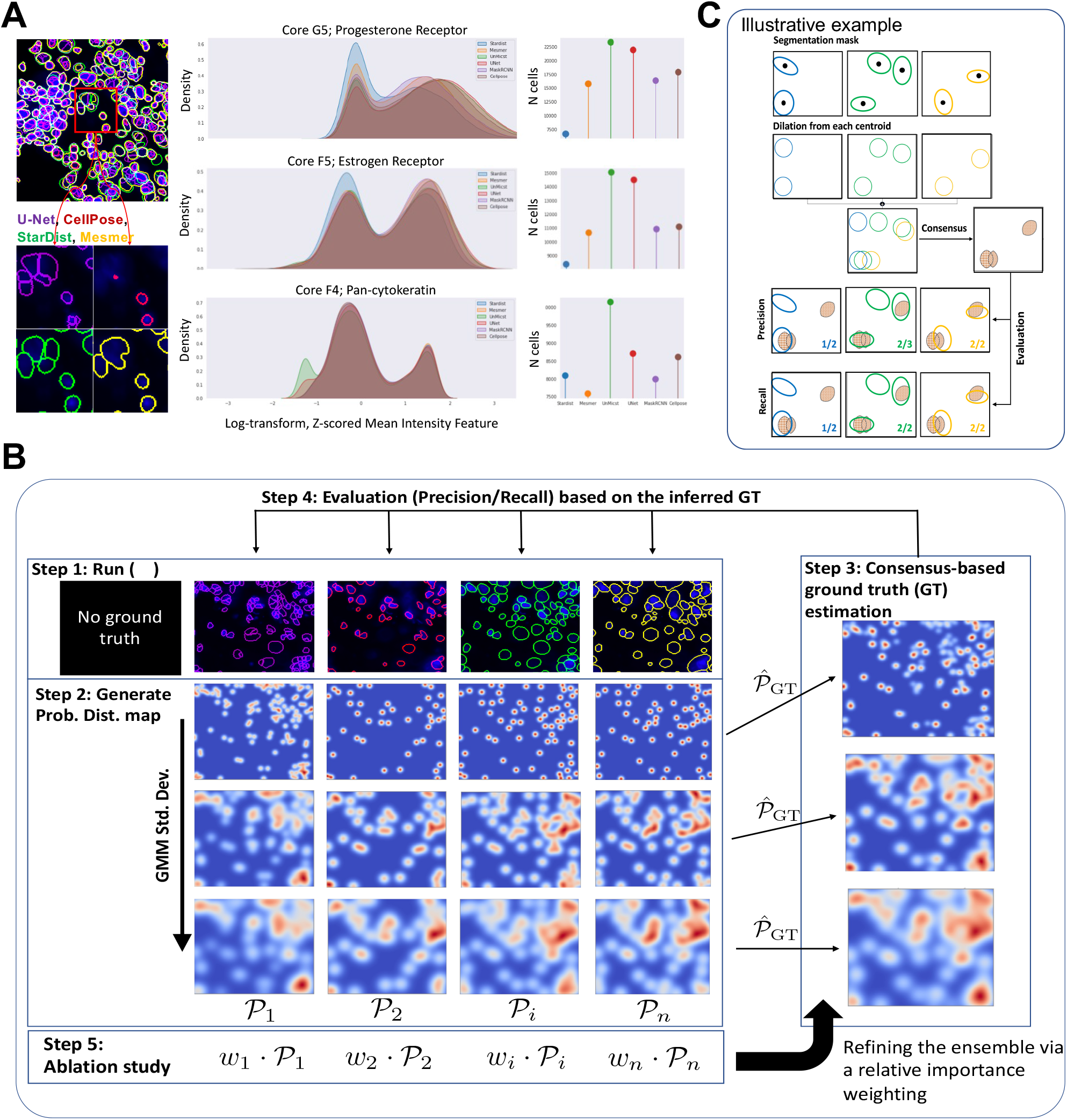
An overview of consensus-based ground truth estimation, refinement, and segmentation evaluation. (A) examples of individual segmentation masks and feature level discrepancy between segmentation methods. (B) workflow of the proposed approach. (C) an illustrative example of ground truth estimation and evaluation.

High-performance cell and nuclei segmentation tools are built with deep neural networks, which are typically trained in a fully supervised or semi-supervised fashion^13^. These algorithms typically rely on extensive training datasets of human-labeled images^8,9,14^ which are laborious to produce and cost or time-prohibitive in most settings. As a result, pre-trained models are often used for biological segmentation on user’s datasets without having ground truth labels. Thus, the evaluation of segmentation performance is based on the user’s visual inspection or requires additional manual annotations to quantitatively evaluate performance. Recent end-to-end pipelines such as MCMICRO^1^ can run multiple segmentation algorithms in parallel, allowing their performance to be compared directly, which is particularly useful and helpful for evaluating segmentation tasks but lacking in the ability to assist non-computational experts in choosing the most appropriate pre-trained model without having ground truth label. Thus, users still rely on their visual inspection with few selected regions of interest or samples, instead of using a systematic approach based on quantitative metrics to score individual performance in the entire dataset.

Thus, challenges arise: how can users objectively evaluate individual segmentation methods for their own dataset without ground truth labels and choose the most suitable segmentation? Previous studies have suggested segmentation evaluation in absence of ground truth labels^15–18^. For example, reverse classification accuracy^15^ uses segmentation results as pseudo-ground truth labels to train a new segmentation model. The accuracy of the “reverse”-trained classifier is used as a proxy for the quality of predicted segmentations (i.e., pseudo ground truth). The primary downside of this method is its reliance on the existence of annotated reference datasets from the same or similar domain as the unlabeled data. A similar approach is provided by training a regression model to predict segmentation error from hand-crafted features of the predicted segmentation mask^16^. As training is fully supervised, it requires several annotated datasets. All the approaches aforementioned require additional model training and are also based on existing labeled datasets from the same domain that may not even exist, making them slow and potentially unreliable. Another class of segmentation evaluation approach is based on a probabilistic, ensemble-based approach to compute precision and recall metrics in the context of a document retrieval task with missing or uncertain ground truth^17^. An ensemble of predictions is known to make better predictions and outperforms any single contributing model^19^. Scores are computed purely probabilistically without thresholding the ensemble average to estimate the ground truth. Recent study^18^ proposed an approach that seeks to evaluate cell segmentation methods by providing an objective evaluation approach based on assumptions about the desired characteristics of good cell segmentation methods. Evaluation metrics rely on the similarity between two segmentation methods and an overall segmentation quality score for each method uses the metrics for all methods with and without perturbation (i.e., added noise and down-sampling). Although this approach does not require reference segmentation, the quality score relies on simple statistics from the pre-defined features.

Here, motivated by the previous work from Lamiroy *et al*.^17^, we developed algorithmic and software tools for evaluating segmentation tasks and selecting from several segmentation methods in absence of ground truth labels. We demonstrate that this approach can infer a pseudo ground truth label based on an ensemble of segmentation results from pre-trained models, and quantitatively evaluate the individual model performance by measuring precision and recall based on the inferred pseudo ground truth label. A framework acknowledges sensitivity to the “collective bias” of the ensemble and thus proposes a weighted ensemble in settings where a relative importance weighting across segmentation methods ameliorates this issue. We first perform these analyses on a small dataset with ground truth annotations to demonstrate the feasibility of the proposed approach. Second, we strengthen the validation by applying the proposed approach on an unlabeled larger tissue microarray (TMA) dataset and provide decision guidelines for the general user to choose the most suitable segmentation methods based on quantitative metrics to score individual performance in the entire dataset.

## RESULTS

### Generating masks and ground truth inference

We propose to estimate a probabilistic ground truth label mask using a majority vote amongst an ensemble of segmentation results from different segmentation methods. Figure 1B illustrates an overview of the proposed approach including consensus-based ground truth estimation, refinement, and segmentation evaluation. Our approximate ground truth can be understood as an ensemble of independent spatial point spread functions (PSF) from the identified centroid of individual cells or nuclei.

To do this, we first started by running all segmentation methods on the entire dataset (Step 1, see METHODS section). Each segmentation method generated nuclear masks (Supplementary Figure 1A), and each method’s mask instances are used to define a single cell object’s probability map so that there is a one-to-one correspondence between segmentation instance centroids and their probability distribution (Step 2). Each centroid of the segmented object from individual segmentation methods is used for generating a two-dimensional Gaussian probability distribution map based on the centroid position. The covariance matrix of each PSF component might be a fixed constant that varies a model hyperparameter or may be set according to instance morphological features. For example, fibroblasts are flat and spindle-shaped so that multivariate asymmetric gaussian distribution would fit better. Herein we simply use a fixed constant for the covariance matrix of each PSF component and this modification would be extended in future work.

Once we have a probability distribution map (*P*_*i*_) from each segmentation method for a given covariance, we estimate pseudo ground truth label masks 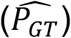 by using a majority vote amongst an ensemble of probability distribution maps from individual segmentation results with the equal weighting factor (Step 3):

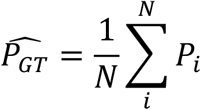

where *N* is the number of segmentation methods. The underlying hypothesis is that when probability distribution maps of individual segmentation model results are correctly combined, we can obtain more accurate and/or robust probability distribution maps of the individual cell/nuclei location. It has been shown that ensemble methods usually aggregate heterogeneous predictions from the different models and produce more accurate solutions than a single model would^19–21^.

### Approximating ground truth pseudo probabilistic label mask and evaluation

In practice, whole-slide MTI data can contain millions of cells/nuclei instances. Modeling, sampling from, and scoring values in such a PSF are computationally expensive. Thus, we simplify our probability model by constructing approximate, binary spatial densities for each segmentation result and taking the average across all methods to generate a pseudo ground truth probabilistic map.

Concretely, for each segmentation we approximate a PSF by removing segmentation boundary information, retaining only an infinitesimally small seed at each cell centroid, and uniformly dilating each seed with a radius *r*. Stacking these binarized masks, summing and normalizing along the channel dimension result in a spatial probabilistic map normalized between 0 and 1. Like the PSF’s covariance matrix, the dilation radius is a model hyperparameter and can be easily replaced by an ellipsoid dilation filter specific to cell instance morphology.

The dilation radius effectively encodes an assumption of how dense the PSF’s probability mass is near the centroid. This assumption follows from the observation that each PSF component should retain most of its probability mass within the segmentation boundary. When increasing the dilation radius in densely populated cells of tissue samples, it is likely that a single segmentation instance from any method ends up composing multiple proxy ground truth objects (see Supplementary Figure 1B). To avoid overcounting, we enforce the additional constraint that each instance can vote for only one ground truth region.

Finally, the approximate pseudo ground truth probability map is binarized such that spatial regions of majority agreement over all segmentation methods are retained and all others are dropped. The result is an approximate pseudo-ground truth against which we evaluate each segmentation method’s precision and recall scores as shown in Figure 1B (Step 4). An illustrative example of evaluation is shown in Figure 1C, where detection is defined as a single-pixel overlap.

### Systematic ablation study refines the ensemble via an adaptive weighting

In Step 3, multiple models are used to infer the pseudo ground truth locations of cells or nuclei. The contribution of each model is considered as a separate vote and the pseudo ground truth inference we get from the majority of the models is used as the final estimation. However, we often observe that certain models might show poor segmentation performance, especially when images under study are very different from the training images of the pre-trained models. To address this, we use systematic model ablations to define the importance of each model for ground truth inference and further improve the ensemble method’s pseudo ground truth inference by assigning refined weights.

To do this, we use the leave-one-out validation method on the ensemble to understand the individual effect of each segmentation method on precision and recall scores. New precision and recall scores are calculated by using the leave-one-out coupled with the calculation of performance change in precision and recall. The change in performance informs us about each specific segmentation method’s positive or negative effect on the ensemble estimation and performance. To calculate the change in the precision of a single method, the average of precision scores available for that method is calculated first. For example, when we compare *N* nuclei segmentation methods and if we consider the *i*-th method, we calculate *N-1* precision scores 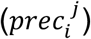 by considering all possible leave-one-out (i.e., *j* = {1, …, *N*} where *j* ≠ *i*), and then take the average of these scores:

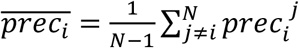

Each precision score of the *i*-th method 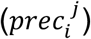 is subtracted from the average value 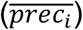, and then the result is divided by the average again to normalize it. For a given radius, the *i*-th method’s precision change can be calculated by

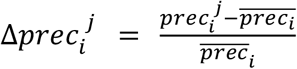

This is repeated for all the methods and for different dilation radii to study the contribution of dilation radius in the estimation. The change in recall 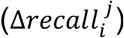 is also computed using the same method for precision change step respectively.

Once the contribution of each method is calculated by using precision and recall score changes, we use a simple average of precision and recall score, and a SoftMax function is applied to calculate method-specific individual weights (*w*_*i*_) as follows:

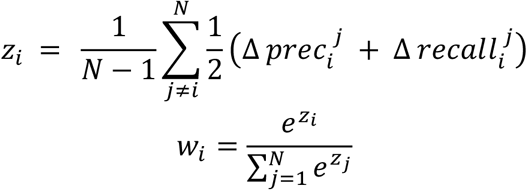

By applying a Softmax or normalized exponential function, the ensemble of weights is normalized (sum equal to 1) and refined weights between methods are applied as shown in Figure 1B (Step 5). For instance, the method that decreases the precision and recall when removed positively affects the ground truth inference. On the other hand, one which increases the precision and recall when removed has a negative effect on the ground truth inference. Finally, the consensus-based ground truth is estimated by using refined weights (see Supplementary Figure 1B and 1C with varying radii), and precision and recall scores are re-evaluated. The method’s overall performance (F1 score) is then calculated as the harmonic mean of the new precision and recall scores.

### Correlation between the proposed metric and Dice coefficient

In this section, we demonstrate the feasibility of the proposed approach to evaluate segmentation performance in 5 tissue microarray (TMA) cores with human ground truth annotations (see details in METHODS section).

We first investigate whether method-specific weighting via ablation study improves the ground truth inference. To test this, we calculate the Dice coefficient – often used to quantify the performance of image segmentation methods - between inferred pseudo-ground truth labels from the ensemble of models and ground truth annotations. Figure 2A shows the Dice coefficient for four commonly used nuclei segmentation methods including Stardist^2^, Cellpose^3^, Mesmer^11^, and UnMICST^12^. The blue bars show results when using equal weights and a dilation radius optimally chosen as 12 pixels (7.5 µm). The inferred pseudo ground truth from the ensemble of segmentation results with equal weights shows a good dice (>0.8) with the ground truth annotations.

**Figure 2.**
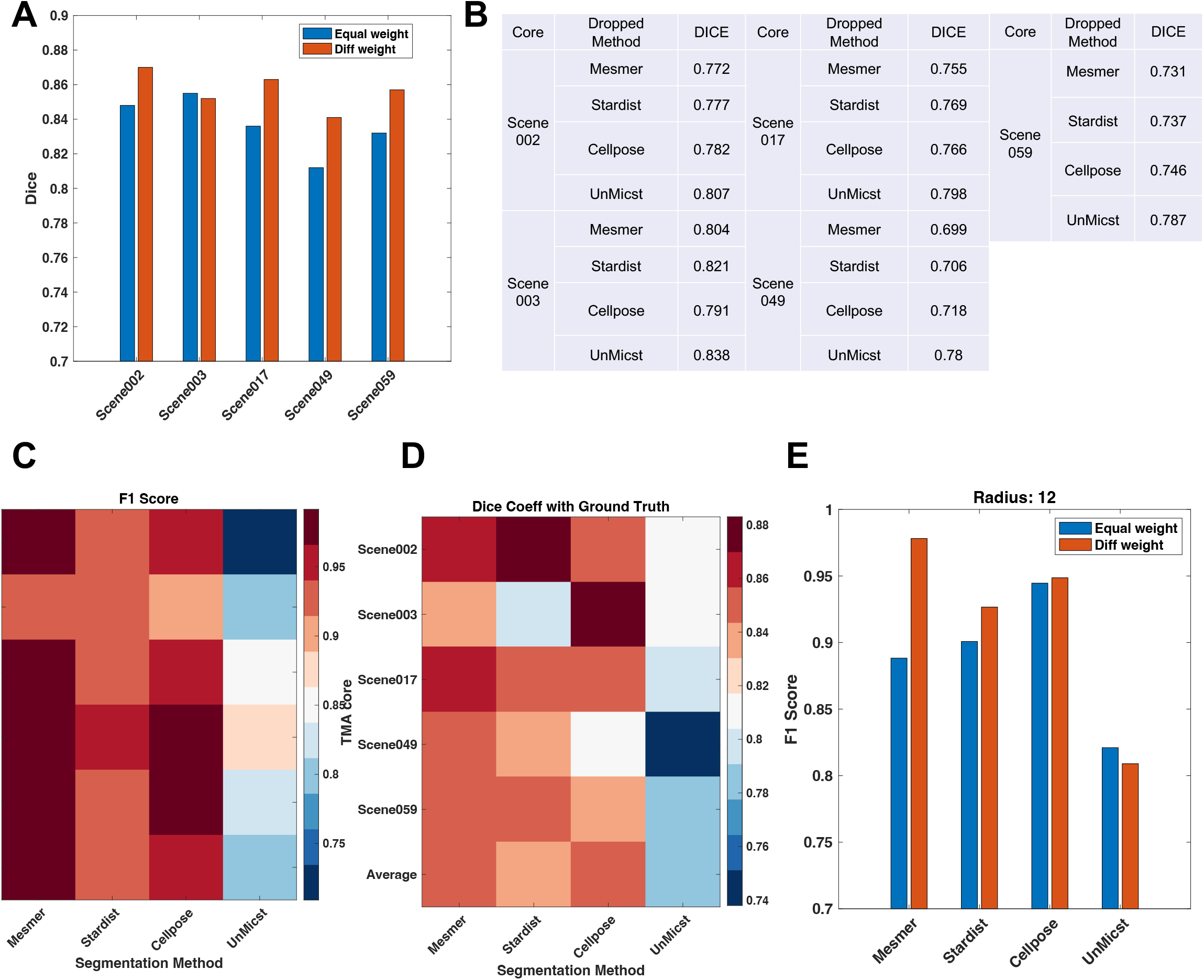
Method-specific weighting via ablation study avoids potential sensitivity to collective bias. (A) Dice coefficient between ground truth label and inferred ground truth with equal weights and different weights respectively. (B) Dice coefficient changes from the leave-one-out segmentation method. (C) F1 score based on the proposed approach with different weights across individual TMA cores. (D) Dice coefficient with ground truth labels across individual TMA cores. (E) averaged F1 score comparison between equal weights and different weights across 5 TMA cores.

Then, we evaluate the Dice coefficient based on the leave-one-out validation method in the ensemble and calculate a relative importance weighting as we proposed. Figure 2B summarizes the Dice coefficient when a specific segmentation method is dropped. The best performer is the method that reduces the Dice coefficient significantly when removed. As an example, for scene 002, when UnMICST is dropped, the overall Dice coefficient increases (i.e., 0.807, *Δ*+0.022) compared to the average Dice coefficient (0.785). On the other hand, if Mesmer is dropped, the overall Dice coefficient decreases (i.e., 0.772, *Δ*-0.013). Based on these observations, we calculate relative importance weights via ablation study and re-evaluate the overall Dice coefficient as shown in Figure 2A (red bars). 4 TMA cores among 5 show improvements for Dice coefficient in the ensemble with refined weights. This suggests that the original inferred pseudo ground truth using the equal weights may be skewed by a poor segmentation method, but with refined weights, the corrected pseudo-ground truth is further closer to the ground truth annotations as shown in Figure 2A.

We next rank individual segmentation methods by computing the overall performance thanks to the proposed approach and with respect to the ground truth annotations. The proposed approach with individually refined weights reports that Mesmer is the best method, followed by Cellpose, Stardist, and UnMICST (Figure 2C), which is consistent with the ground truth-based ranks (Figure 2D). Also, the overall performance trend of each TMA core matches with the results using the Dice coefficient with ground truth. Note that we use F1 score to measure overall performance for the proposed approach, and precision and recall scores are shown in Supplementary Figure 2A; overall, all segmentation methods show good precision scores (>0.9), but with variable recall scores. Thus, the recall score is the most important metric when performing an F1 score-based classification, and this trend is consistent with the Dice coefficients observed in Figure 2D.

In Figure 2E, we also compare the F1 scores obtained with equal and refined weights. With equal weight, Mesmer is ranked 3^rd^, but by refining the ensemble with weighting factors, it is ranked as the best performer. Thus, it is important to evaluate individual model contributions to the ensemble and refine the ensemble by using adaptive weighting. We show that the performance measured by the proposed approach is consistent with the Dice based on the ground truth annotation. We also report the performance with varying dilation radii (*r* =12, 16, and 20). Supplementary Figure 2B shows that the overall trend is similar, but as the dilation radius increases, the overall F1 score decreases slightly. It is likely because a single segmentation method ends up composing multiple proxy ground truth objects as the dilation radius increases as shown in Supplementary Figures 1B and 1C.

### Performance comparison and evaluation without ground truth label

We further evaluate the proposed approach in a large TMA dataset without ground truth. We first determine the relative importance of weights via ablation study with varying dilation radii. Figure 3A shows performance change (i.e., F1 score) in the ablation study. Overall, we observe that U-Net^22^, and UnMICST show a negative effect on the performance, and Mesmer, Stardist, Cellpose show a positive effect on the overall performance similar to Mask R-CNN. Note that Mask R-CNN model was originally trained using 5 manually annotated TMA cores in the BC TMA dataset, so the performance of Mask R-CNN could be used as the reference for performance comparison (see details in METHODS section).

**Figure 3.**
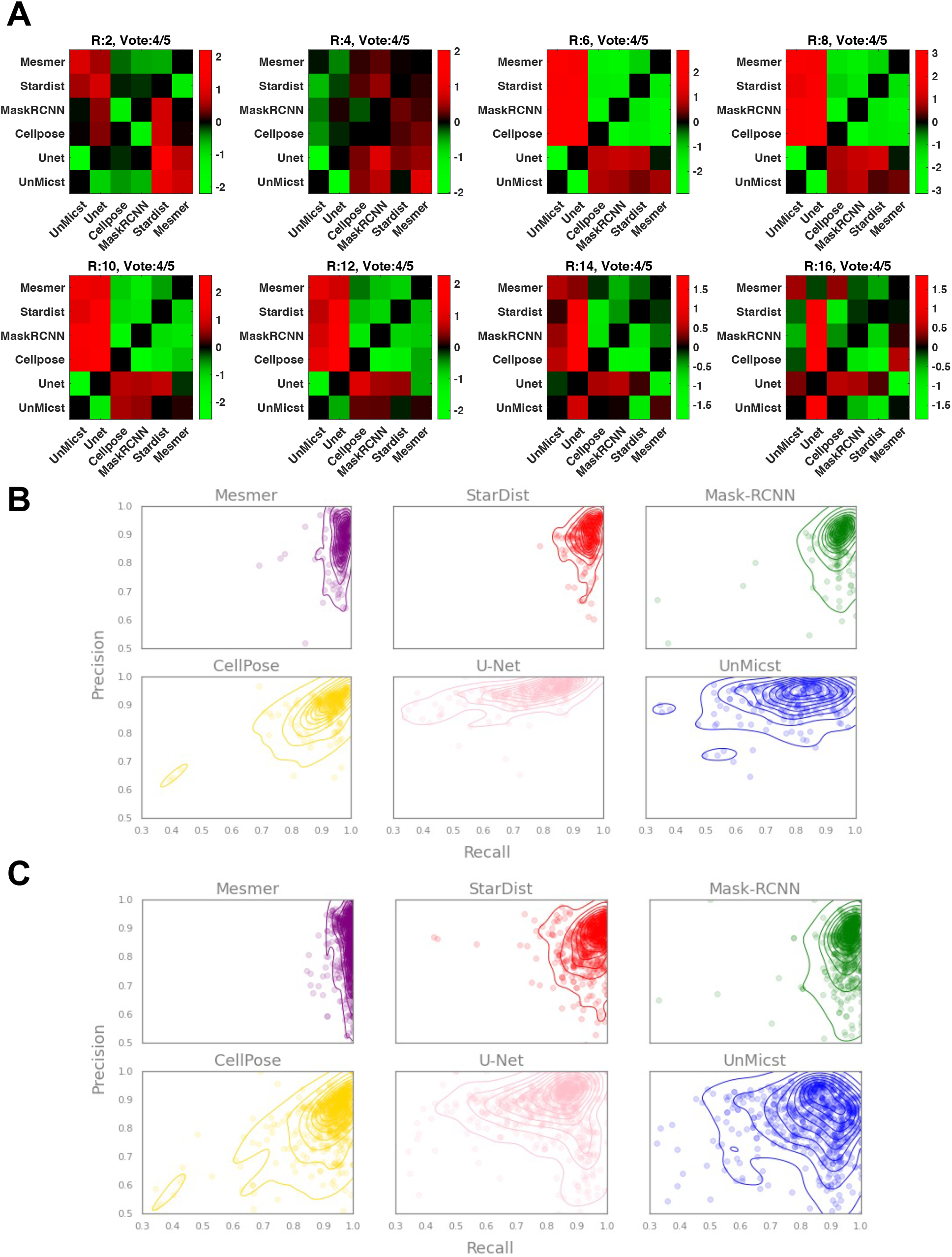
An example of a large TNP-TMA dataset with 6 different segmentation approaches. (A) ablation study determines method-specific weights. (B) precision and recall with equal weights and (C) precision and recall with different weights where an individual dot represents a single TMA core and the optimal radius is selected.

With individually tuned weights via ablation study, Mesmer shows a relatively high recall and good precision score compared to Mask R-CNN performance. Also, we observe that with equal weights, U-Net and UnMICST show good precision scores but performance decreases with different weights (Figure 3B and 3C). This result is consistent with the performance change in the ablation study. Also, we observe that Mesmer shows a small variation of performance (i.e., precision and recall scores) across individual TMA cores compared with Cellpose, U-Net, and UnMICST. This indicates that Mesmer is robust to potential variations in cell types and staining of all cells across BC subtypes. Thus, for the BC TNP-TMA dataset, Mesmer would be the most suitable segmentation method among other methods we used in this study.

In summary, the proposed approach provides accurate performance metrics across individual segmentation methods by inferring pseudo ground truth to help users select the most suitable segmentation methods for their own dataset without ground truth annotation.

## DISCUSSION

We often observe large differences in segmentation performance between different pre-trained deep learning models^23^, which is also true between human annotators^7,24^. Therefore, the variety of segmentation approaches cannot be captured by a single generalizable model although many recent deep learning models claim a generalist algorithm for segmentation^7^.

This variability reflects fundamental aspects of biomedical imaging data, human annotation tasks, and segmentation models. Here, we have addressed how to ensure accurate segmentation among many existing segmentation models and evaluate performance especially when ground truth annotation is not available. We use a probabilistic, ensemble-based approach to capture the variety of biological segmentation styles and infer a pseudo ground truth from the different model segmentation results as there often exist multiple acceptable solutions^23^.

As models in the ensemble exhibit performance discrepancies, we use a weighted ensemble to give individually tuned contributions depending on the inferred ground truth. A systematic ablation study identifies each segmentation model’s contribution to performance change and refines the pseudo ground truth inference. We demonstrate that the ensemble via relative importance weights improves the ground truth inference and corrects the segmentation performance evaluation.

Recent studies have suggested interactive approaches such as human-in-the-loop to retrain the imperfect model with their own annotations in the user’s dataset^23,25^. Future research can look into the deployment of the proposed approach in the human-in-the-loop and iterative annotation and model re-training settings. For instance, the proposed approach can be extended to quantify the quality of additional annotation by scoring individual annotator contributions as well as the refined model evaluation in the ensemble setting. Finally, here we simply use circular dilation, but our future effort will be considering a probability distance map or an ellipsoidal dilation specific to cell instance morphology for further improvement.

## METHODS

### Segmentation models

Here we test U-Net^22^, Stardist^2^, Cellpose^3^, Mesmer^11^, UnMICST^12^, and Mask R-CNN models for inferring ensemble-based ground truth labels and self-evaluating their performance. All methods were used by their default set-up with proper physical resolution information for conversion. For U-Net, a segmentation mask has been performed previously^1^ and thus we simply used the previously generated mask. For UnMICST, we used S3 segmenter (https://github.com/HMS-IDAC/S3segmenter) by default.

Mask R-CNN^26^ is well known and widely used instance segmentation architecture. We used the model recommended and provided in the cycIFAAP pipeline (https://www.thibault.biz/Research/cycIFAAP/cycIFAAP.html). This model is a Mask R-CNN TorchVision implementation, and it was trained with 5 breast cancer (BC) HER2+ tissue microarray (TMA) cores (described in the Dataset section), using pseudo-randomly selected 512×512 crops with a high probability to keep non-empty crops coupled with a simple classical data augmentation (flipping and rotation). This model achieves 96.9% accuracy and 0.83 dice score on the training dataset using leave-one-out cross-validation.

### Datasets

#### 5 selected TMA cores

This dataset is composed of 5 BC HER2+ TMA cores, containing an approximative total of 70K manually segmented nuclei. These images were manually segmented using Microscopy Image Browser (MIB) by a biomedical imaging expert.

#### TNP-TMA dataset

The dataset used for evaluating segmentation methods is a BC TMA available on synapse from the Human Tumor Atlas Network (HTAN) TNP-TMA (https://www.synapse.org/#!Synapse:syn22041595). This BC TMA dataset is comprised of 88 cores and 6 different cancer subtypes: luminal A, luminal B, luminal B/HER+, Triple Negative, and Invasive Lobular Carcinoma.

## Code availability

All software used in this manuscript is detailed in the article’s Results and Methods section and the associated scripts are freely available via GitHub as described at https://github.com/lstrgar/seg-eval. We provide an example Jupyter notebook to reproduce the analyses of Supplementary Figure 1 at: https://github.com/lstrgar/segeval/blob/main/Segmentation Evaluation Methods Breakdown.ipynb

## ACKNOWLEDGEMENTS

We thank Luke Ternes, Eun Na Kim, and Bipasa Bose for helping in all stages of this project. We also thank Jia-Ren Lin and Yu-An Chen (Harvard Medical School) for their help and for sharing the dataset. This work was partly supported by NIH grants (U54CA209988, U2CCA233280, R01 CA253860) and Kuni Foundation Imagination Grants.

## SUPPLEMENTARY INFORMATION

**Supplement Figure 1.**
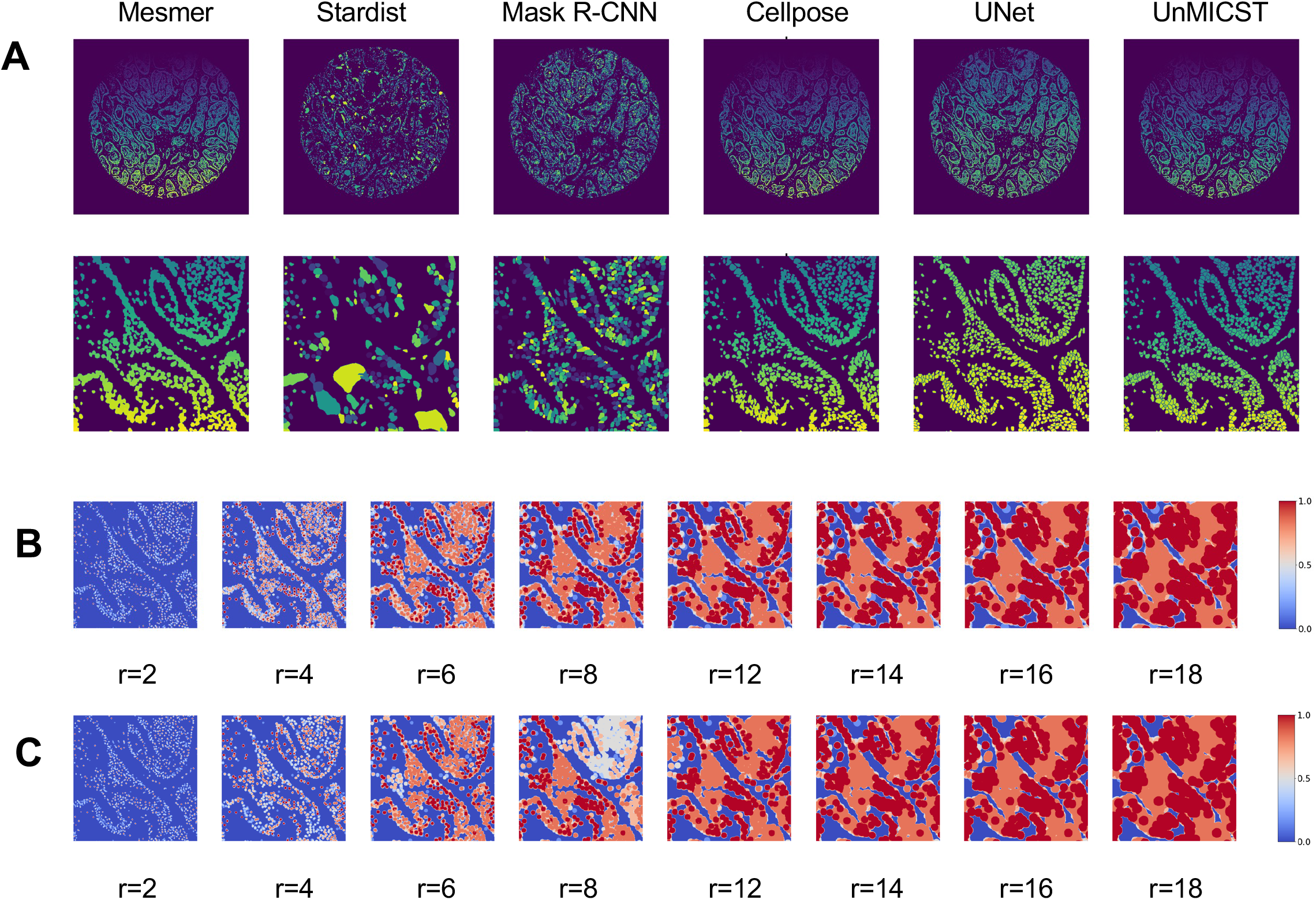
Example images of segmentation masks and resulting pseudo ground truth probability maps. (A) both the entire and zoomed-in sections of segmentation masks from the BC TMA dataset for each method. (B) binarized pseudo-probability maps derived from the masks for different-sized radii (unit: pixels) (C) refined pseudo ground truth probability masks by only counting pixels where there is agreeance among at least 3 of the different methods from (B).

**Supplement Figure 2.**
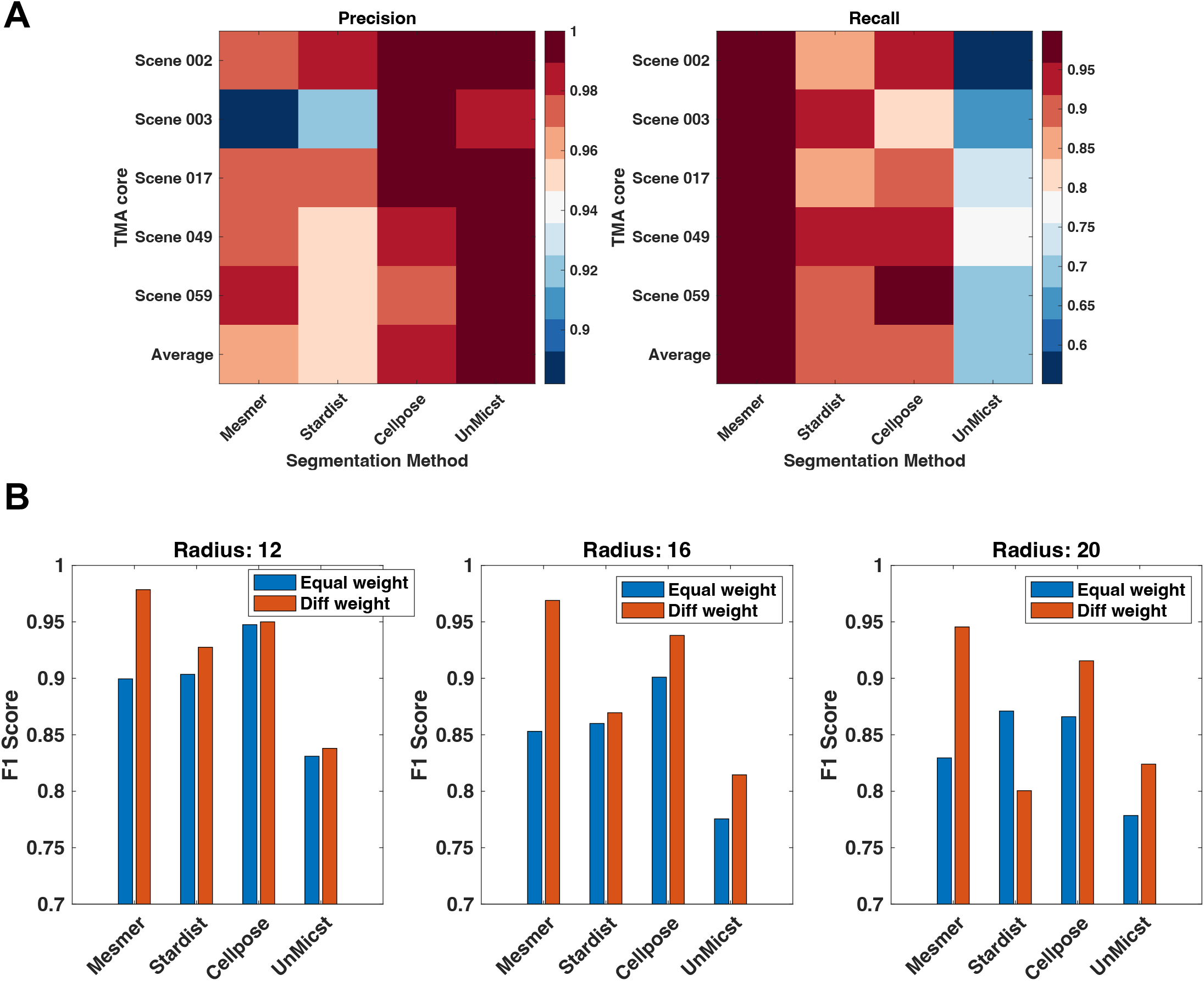
Refined ensemble-derived scores. (A) precision and recall score in 5 TMA cores. (B) F1 score with varying dilation radii.

## REFERENCE

1. Schapiro, D. et al. MCMICRO: a scalable, modular image-processing pipeline for multiplexed tissue imaging. Nat. Methods 19, 311–315 (2022).

2. Tsujikawa, T. et al. Quantitative Multiplex Immunohistochemistry Reveals Myeloid-Inflamed Tumor-Immune Complexity Associated with Poor Prognosis. Cell Rep. 19, 203–217 (2017).

3. Goltsev, Y. et al. Deep Profiling of Mouse Splenic Architecture with CODEX Multiplexed Imaging. Cell 174, 968-981.e15 (2018).

4. Lin, J.-R. et al. Highly multiplexed immunofluorescence imaging of human tissues and tumors using t-CyCIF and conventional optical microscopes. Elife 7, (2018).

5. Burlingame, E. A. et al. Toward reproducible, scalable, and robust data analysis across multiplex tissue imaging platforms. Cell Rep Methods 1, (2021).

6. Schmidt, U., Weigert, M., Broaddus, C. & Myers, G. Cell Detection with Star-Convex Polygons. in Medical Image Computing and Computer Assisted Intervention –MICCAI 2018 265–273 (Springer International Publishing, 2018).

7. Stringer, C., Wang, T., Michaelos, M. & Pachitariu, M. Cellpose: a generalist algorithm for cellular segmentation. Nat. Methods 18, 100–106 (2021).

8. Edlund, C. et al. LIVECell-A large-scale dataset for label-free live cell segmentation. Nat. Methods 18, 1038–1045 (2021).

9. Bannon, D. et al. DeepCell Kiosk: scaling deep learning–enabled cellular image analysis with Kubernetes. Nature Methods vol. 18 43–45 Preprint at https://doi.org/10.1038/s41592-020-01023-0 (2021).

10. Lucas, A. M. et al. Open-source deep-learning software for bioimage segmentation. Mol. Biol. Cell 32, 823–829 (2021).

11. Greenwald, N. F., Miller, G., Moen, E., Kong, A. & Kagel, A. Whole-cell segmentation of tissue images with human-level performance using large-scale data annotation and deep learning. Nature biotechnology, 40(4), pp.555–565 (2022).

12. Yapp, C., Novikov, E., Jang, W. D., Chen, Y. A. & Cicconet, M. UnMICST: Deep learning with real augmentation for robust segmentation of highly multiplexed images of human tissues. Communications Biology, 5(1), p.1263 (2022).

13. Robitaille, M. C., Byers, J. M., Christodoulides, J. A. & Raphael, M. P. Self-supervised machine learning for live cell imagery segmentation. Commun. Biol. 5, 1162 (2022).

14. Caicedo, J. C. et al. Nucleus segmentation across imaging experiments: the 2018 Data Science Bowl. Nat. Methods 16, 1247–1253 (2019).

15. Valindria, V. V. et al. Reverse classification accuracy: Predicting segmentation performance in the absence of ground truth. arXiv [cs.CV] (2017).

16. Kohlberger, T., Singh, V., Alvino, C., Bahlmann, C. & Grady, L. Evaluating segmentation error without ground truth. Med. Image Comput. Comput. Assist. Interv. 15, 528–536 (2012).

17. Lamiroy, B. & Sun, T. Computing precision and recall with missing or uncertain ground truth. in Graphics Recognition. New Trends and Challenges 149–162 (Springer Berlin Heidelberg, 2013).

18. Chen, H. & Murphy, R. F. Evaluation of cell segmentation methods without reference segmentations. Mol. Biol. Cell mbcE22080364 (2022).

19. Bousselham, W. et al. Efficient Self-Ensemble for Semantic Segmentation. arXiv [cs.CV] (2021).

20. Arsov, N., Pavlovski, M., Basnarkov, L. & Kocarev, L. Generating highly accurate prediction hypotheses through collaborative ensemble learning. Sci. Rep. 7, 44649 (2017).

21. Wan, Q. & Pal, R. An ensemble based top performing approach for NCI-DREAM drug sensitivity prediction challenge. PLoS One 9, e101183 (2014).

22. Ronneberger, O., Fischer, P. & Brox, T. U-Net: Convolutional Networks for Biomedical Image Segmentation. in Lecture Notes in Computer Science 234–241 (Springer International Publishing, 2015).

23. Pachitariu, M. & Stringer, C. Cellpose 2.0: how to train your own model. Nat. Methods 19, 1634–1641 (2022).

24. Tsujikawa, T. et al. Robust Cell Detection and Segmentation for Image Cytometry Reveal Th17 Cell Heterogeneity. Cytometry A 95, 389–398 (2019).

25. Berg, S. et al. Ilastik: Interactive machine learning for (bio)image analysis. Nat. Methods 16, 1226–1232 (2019).

26. He, K., Gkioxari, G., Dollár, P. & Girshick, R. Mask R-CNN. arXiv [cs.CV] (2017).

